# Identifying protein subcellular localisation in scientific literature using bidirectional deep recurrent neural network

**DOI:** 10.1101/2020.09.09.290577

**Authors:** Rakesh David, Rhys-Joshua D. Menezes, Jan De Klerk, Ian R. Castleden, Cornelia M. Hooper, Gustavo Carneiro, Matthew Gilliham

**Affiliations:** School of Agriculture, Food and Wine and The Waite Research Institute, Waite Campus, The University of Adelaide, South Australia, Australia; ARC Centre of Excellence in Plant Energy Biology, Waite Campus, The University of Adelaide, South Australia, Australia; School of Computer Science, Australian Institute for Machine Learning, The University of Adelaide, South Australia, Australia; ARC Centre of Excellence in Plant Energy Biology, The University of Western Australia, WA, Australia

**Keywords:** Natural Language Processing, Relation Extraction, subcellular localisation, text mining

## Abstract

With the advent of increased diversity and scale of molecular data, there has been a growing appreciation for the applications of machine learning and statistical methodologies to gain new biological insights. An important step in achieving this aim is the Relation Extraction process which specifies if an interaction exists between two or more biological entities in a published study. Here, we employed natural-language processing (CBOW) and deep Recurrent Neural Network (bi-directional LSTM) to predict relations between biological entities that describe protein subcellular localisation in plants. We applied our system to 1700 published *Arabidopsis* protein subcellular studies from the SUBA manually curated dataset. The system was able to extract relevant text and the classifier predicted interactions between protein name, subcellular localisation and experimental methodology. It obtained a final precision, recall rate, accuracy and F1 scores of 0.951, 0.828, 0.893 and 0.884 respectively. The classifier was subsequently tested on a similar problem in crop species (CropPAL) and demonstrated a comparable accuracy measure (0.897). Consequently, our approach can be used to extract protein functional features from unstructured text in the literature with high accuracy. The developed system will improve dissemination or protein functional data to the scientific community and unlock the potential of big data text analytics for generating new hypotheses from diverse datasets.

## Introduction

Experimental techniques to characterize proteins at a biochemical, structural and physiological level have improved considerably in the last 25 years providing researchers with the tools necessary to understand protein function at a cellular and an organism level. Combined with detailed functional data, large-scale genome sequencing efforts have also greatly increased the scale of proteomic data available from model and non-model species. However, a major challenge facing researchers today is simply keeping pace with the sheer volume of low-throughput and high-throughput data being generated. Although scientific publications are used to disseminate research findings to the wider community, manually identifying, curating and collating individual experiments is a time and labour intense process. In addition, the large number of papers published, and the unstructured and versatile format of the content make the data difficult to integrate and analyse. The development of biological databases to curate protein experimental data from published literature have, as a result, become indispensable research tools for researchers. However, manual curation makes it difficult for databases to keep up with the ever-increasing amounts of data being generated.

To address these issues, automated and augmented curation systems for extracting protein functional data from scientific literature is becoming increasingly desired. In particular, Machine Learning and Natural Language Processing techniques are beginning to be employed for large scale biocuration efforts^1,2^. Biocuration refers to the process of extracting and organising unstructured biological information into a structured form that is accessible to biologists. Central to these automated systems, is the process of unambiguously extracting semantic relationships between two or more biological entities in the literature^3^. When referring to proteins or genes, entity relationships can describe protein-protein interaction, drug interactions, physicochemical properties or functional motifs, disease/trait interaction, and are very useful for biological network construction^4–7^. In addition, ML techniques have the potential to assist in protein annotation efforts such as the manually curated UniProt database by automatically extracting protein knowledge from research articles^2^.

Initial methods for entity relationship extractions (RE) relied on dictionary-based and rule-based methods and parsers^8^. This type of system worked well for entity detection if the dictionary was big but still struggled to extract relations especially if there was a large distance between related words. Further improvements to biological entity RE were described by Fundel et al. (2007) using a multi-step process^9^. The approach relies on first preprocessing data and then creating a parse tree of the sentence that can be filtered against extraction rules. Other methods rely on Support Vector Machines (SVM) and kernel-based solutions or by combining the two approaches^10^. In the combined approach, kernels were utilized by converting the input text to a vector format and then finding the similarity between two entities and the sum of their sub structures similar to a “bag of words” approach. A SVM was then used for entity classification. This gave both the benefits of kernels such as being able to search a large feature space and regularization methods such as boundary detection between classes from SVM’s. These types of methods improved the task of relation classification but ultimately were suboptimal in the field of relation detection.

With the advent of machine learning, many methods started to treat words as sequence data and applied deep learning models to find solutions. In the study by F. Li et al. (2017) the authors suggest creating a dependency tree which then feeds into a Convolutional Neural Network (CNN) to pretrain character level embeddings from words in the tree^11^. The tree was then passed into a bi-directional Long Short-Term Memory (LSTM) model with the word embeddings and eventually to a dense layer. The end result was then put through a softmax layer and returned a binary classification corresponding to if the sentence contains an entity relation or not. It is shown that this method gets good results, but a major disadvantage is that due to the dependency tree, inter-sentence relations are not able to be extracted, limiting the scope of the biological entity integration.

Furthermore, text analytic methods described for biological research are largely optimised for extracting binary entity relationships such as interaction between two proteins or association between proteins and diseases^12^. Often, the experimental methodology that was used to verify the result, although mentioned in the sentence, is not processed as an entity type and hence not linked to the protein. The experimental methodology is an important factor for interpreting the results. For example, when analysing protein subcellular localisation, two commonly used approaches are fluorophore tagging of the protein (e.g. Green Fluorescent Protein) or by a proteomic approach of identifying all the proteins within a specific cellular compartment (e.g. Mass Spectrometry). Each approach has their advantages as the fluorophore approach provides better spatial resolution of protein location whereas MS methods allow for better quantitative analysis of cellular proteomes. Knowing what technique was used together with protein subcellular information can guide researchers in designing future experiments.

In this study, we describe the development and implementation of a pipeline to extract protein subcellular information and the experimental methodology from published studies using deep learning techniques. Protein subcellular location was chosen as a feature to extract as knowing where a protein resides within a cell provides important clues to its cellular function and represents a fundamental unit of how proteins function in nature. We tested the deep learning system across two datasets, one for papers describing protein subcellular localisation in Arabidopsis (SUBA)^13^ and another dataset that includes four major crop species (CropPAL)^14^. We demonstrate, through this method, entity relationship can be predicted with high accuracy. Thus, the pipeline provides a framework for high-throughput extraction and linking of biological entities from unstructured text giving researchers access to the latest scientific information from diverse datasets.

## Results and Discussion

### Outline of the approach

We describe a semi-automated pipeline for extraction of complex protein features from unstructured published literature (Fig. 1). The process includes text parsing to identify sentences within an article that contain relevant biological entities, followed by extraction of relationships between entities. In order to test the validity of our approach, we analysed articles describing subcellular localisation of proteins experimentally determined by fluorescence tagging and mass spectrometry, two commonly used techniques to resolve spatial and quantitative properties of protein in living cells^13^. Three biological entity types were considered for this classification: protein name, experimental methodology and subcellular location. The deep learning model was assessed for its ability to classify ‘True’ or ‘False’ relationships between these three entity types. A True relationship indicates the given protein was experimentally verified to be located within a specific cellular organelle and a False relationship indicates the co-occurrence of these entity types without being related. Implementation details and source codes of the pipeline are available at GitHub: https://github.com/RhysMenezes/find-a-protein

**Figure 1.**
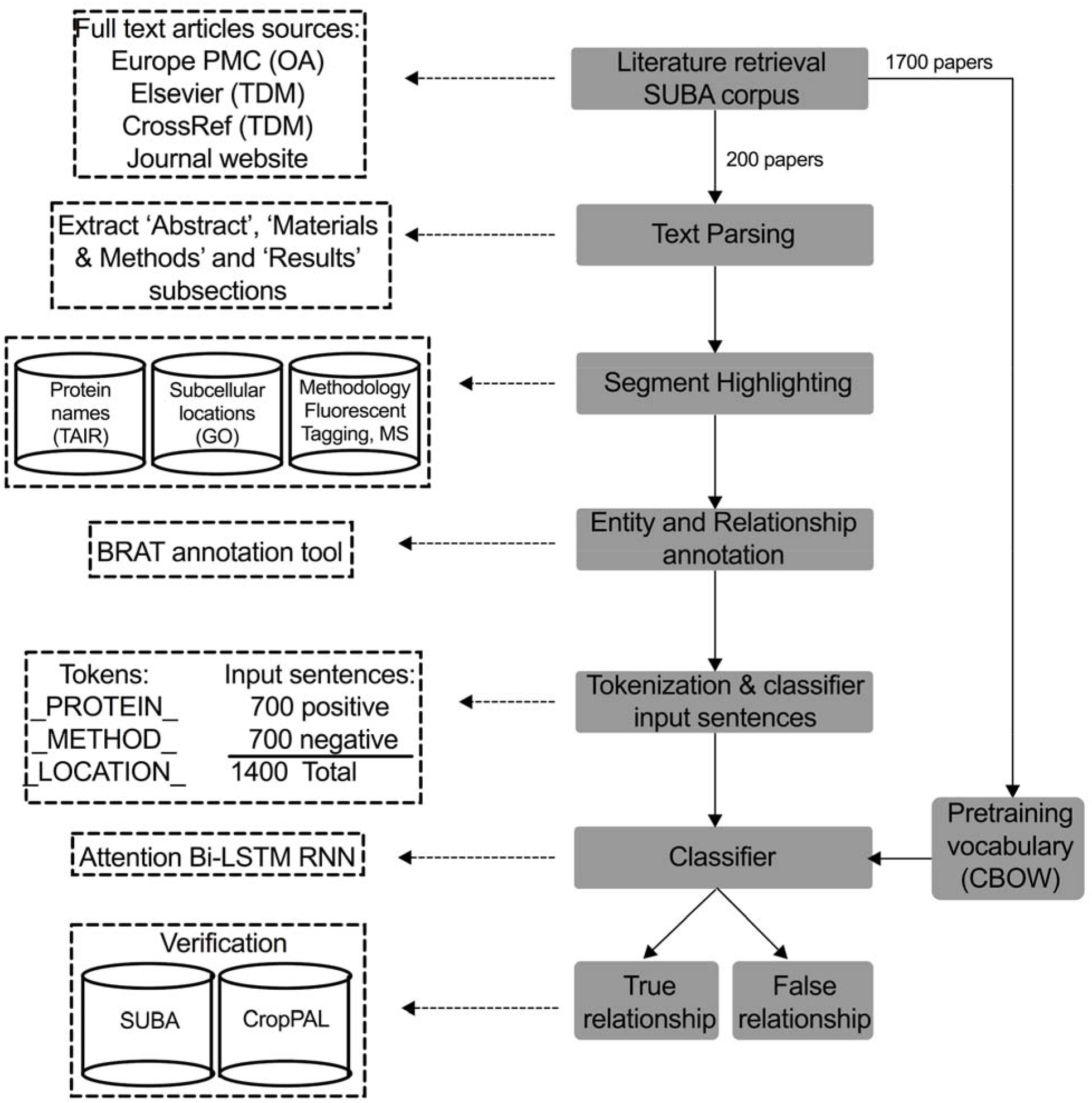
Overview of the deep learning approach to extract protein subcellular information from scientific literature. A summarised workflow of the system is shown on the right (grey boxes). Technical details for the implementation of each step including data sources, methodology and input examples is shown on the left (dashed boxes). The system was trained and tested using published studies from the SUBA corpus of protein subcellular data in *Arabidopsis*. Using the Pubmed ID in SUBA, full text articles were bulk retrieved from Europe PMC open access (OA) repository, text and data mining (TDM) services from Elsevier and CrossRef as well as directly from journal publisher website. A subset of 200 articles were used for annotating protein subcellular information to be used subsequently for classifier training and testing purposes. Triplet entities and relationships were annotated using the BRAT tool, following which protein names, experimental methodology and subcellular location were replaced with tokens to create input sentences for the RNN classifier. A second neural network, Continuous Bag of Words (CBOW) was employed to pretrain the classifier using the complete 1700 full-text articles from SUBA. The classifier determines a binary true or false classification for an input sentence to contain a valid triplet entity relationship. The results of the classifier were manually verified using Pubmed ID as queries in the SUBA online database and transfer learning ability of predictive model was demonstrated using an independent dataset from the CropPAL corpus.

### Dataset and literature retrieval

The SUBA collection (version 4.0) (http://suba.live/) includes subcellular information relating to 11,740 proteins that was manually extracted by expert curators from 1,768 published studies and stored in a MySQL format together with full citation details^13^. Full-text was required for the analysis, as often the protein subcellular information, and particularly the supporting experimental evidence are not described in the abstracts accessible in PubMed. Using a combination of strategies including an in-house developed NLP publication retrieval and processing tool outlined in the Materials and Methods, we were able to analyse 1700 full-text articles for protein subcellular information.

### Parsing and segment highlighting

Published full-text articles were processed into XML-tagged documents, and the ‘Abstract’, ‘Results’ (including figure captions) and ‘Materials and Methods’ subsections were extracted as these are likely to include protein subcellular information and experimental methods. Given the time-consuming process of manual annotation, we chose a random subset of 200 studies from the SUBA corpus to annotate entities and relationships. To identify relevant text, article subsections were parsed using regular expressions terms to find co-occurrences of three biological entities; protein names, subcellular locations and experimental methodology in a forty-word sliding window. Entity names were matched against a list of all Arabidopsis protein names (including synonyms) from The Arabidopsis Information Resource (TAIR), subcellular compartment names derived from the GO cellular components annotations^15^ and fluorescence tagging methodologies (Supp table S1-S2). Using this strategy, passages of text that include one or more triplet entity groups as well as neighbouring contextual words were extracted and used as the primary data for testing and training the recursive classifier.

### Annotation of entity types and relationships

From the 200 pre-processed papers, we retrieved 1400 sentences from the segment highlighting approach and these were subsequently annotated for one or more triplet entity groups. In order to validate the correct triplet entity was being used for the deep learning prediction model, entities were labelled and the relationship between them were specified using the BRAT manual annotation tool^16^ (Fig. 2, Supplemental Fig. S1). Protein name was used as the common entity type to specify the relationship between the methodology and subcellular location described in the text. As the description of protein names can vary in individual studies, three entity types were used for annotation; abbreviated names of proteins including synonyms (‘protein’) (TAIR), modified proteins as result of mutations, truncations (‘protein_variant’) and proteins that are described as fluorescent tagged versions (‘tagged_protein’). With relation to subcellular studies, we found tagged proteins in which both a protein name and methodology is concatenated in the same word, (e.g. SUVR2a-GFP^17^), were the most commonly used protein descriptors in the SUBA corpus. In such cases, the concatenated word gets labelled as one entity type called ‘tagged protein’ and sub labels are used to differentiate between protein and methodology (Supplemental Fig. S1, Supplemental table S3)^18,19^.

**Figure 2.**
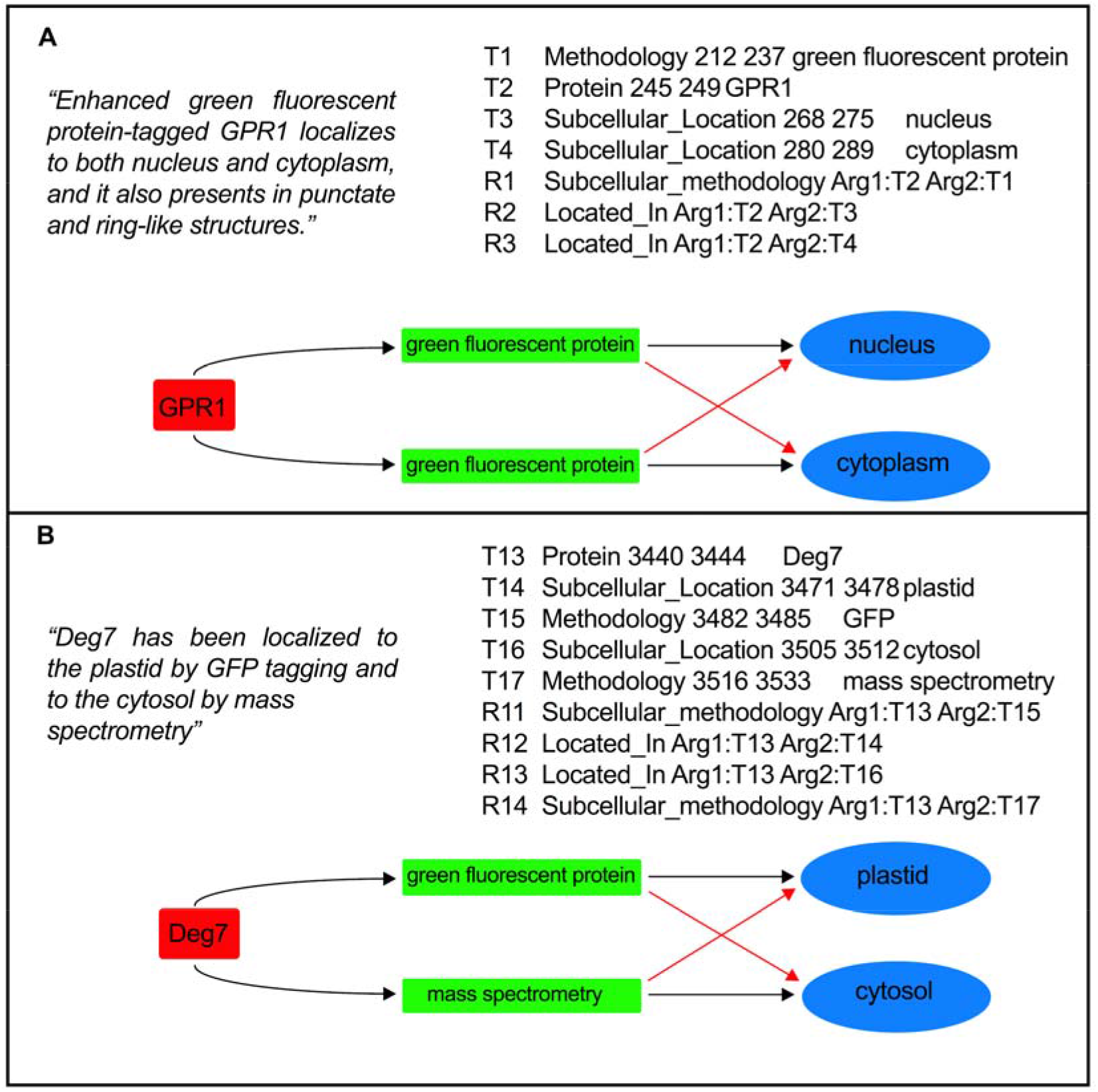
Two examples of protein subcellular information described in the SUBA corpus text and mapping of entity relationships. For both examples, the sentence from the article is quoted (top left), BRAT annotation defining the entities and relationships for that sentence in standoff format (top right) and a visual network representing the interaction between protein (red), experimental methodology (green) and cellular location (blue) is shown (bottom). The example shown in panel A is from Yang et al. and illustrates a simple semantic and syntactic relationship between the protein, GPR1, experimental methodology using green fluorescent protein (GFP) and subcellular locations, nucleus and cytoplasm^20^. In this example, the classifier predicting the relationship for each triplet entity group is straightforward as both groups use GFP as the common methodology. In this case, an incorrect classifier prediction between the two entity groups (red arrows in both panels) would still result in an overall correct relationship being defined. The example shown in panel B is from Tanz et al. (2014) and illustrates a more complex sentence in which Deg7 protein is demonstrated to be located in the plastid and cytosol using GFP and mass spectrometry, respectively^21^. Here, the classifier would need to use the contextual words in the input sentence to correctly identify the syntactic relationship between the three entities in each group.

### Annotation cleaning and tokenization

The RNN model’s task is to classify an input sentence as either “True” relationship or “False” relationship, the output is discrete either a 1 (true) or a 0 (false). For all input sentences annotated, a “True” relationship is defined as one that satisfies both semantic and syntactic relationship between a triplet entity group. The entities are considered to have a semantic relationship if they are structured such that the protein’s subcellular location and the methodology used can be logically discerned from the sentence (Fig. 2)

For the input data used in the RNN classifier, the actual protein names, methodologies and subcellular locations were not needed as we investigated the possibility of classifying relationships using the entities neighbouring context words. The names in the input sentences were replaced by tokens that represent the entity types. All protein names were replaced with a “_PROTEIN_” taken, methodologies were replaced with a “_METHOD_” token and subcellular locations were replaced with a “_LOCATION_” token. As the model classifies if a given relationship is true or false, the input sentences will have to be presented such that only one relationship is present in a sentence. If a sentence contains multiple tokens then a valid triplet will be tokenized, and the rest blanked out using _BLANK_ tokens. For example, the sentence from Tanz et al., (2014) “Both Deg1 and Deg9 have been localized to the plastid and nucleus using mass spectrometry”^21^, four separate true states, one for each triplet group (underlined) can be presented to the classifier:

1. “Both ***_<u>PROTEIN</u>_*** and ***_BLANKP_*** have been localized to the ***_<u>LOCATION</u>_*** and ***_BLANKL_*** using mass ***_<u>METHOD</u>_***.”
2. “Both ***_<u>PROTEIN</u>_*** and ***_BLANKP_*** have been localized to the ***_BLANKL_*** and ***_<u>LOCATION</u>_*** using ***_<u>METHOD</u>_***.”
3. “Both ***_BLANKP_*** and ***_<u>PROTEIN</u>_*** have been localized to the ***_<u>LOCATION</u>_*** and ***_BLANKL_*** using ***_<u>METHOD</u>_***.”
4. “Both ***_BLANKP_*** and ***_<u>PROTEIN</u>_*** have been localized to the ***_BLANKL_*** and ***_<u>LOCATION</u>_*** using ***_<u>METHOD</u>_***.”

Lastly to ensure all input data was consistent and the dimensions remained the same in the classifier model, sentences were padded using the PAD term to the maximum sequence length. From 1400 tokenized text input sentences, we created 700 positive examples that satisfied both semantic and syntactic criteria for entity relationships. To generate negative cases, we randomly permutated the entities within the sentences to create 700 negative examples.

### Pretraining - CBOW

In order to utilize the SUBA data more effectively and due to the relatively small number of annotated sentences used for training the RNN classifier, we pretrained the network’s vocabulary on the complete SUBA dataset comprising of 1700 full-text papers. Pretraining the model with plant biology specific domain knowledge has the advantage of improving the performance of the RNN classifier in entity relationship prediction. Pretraining was achieved using CBOW NLP model that represents the meaning of each word in a sentence as a single weighted vector representations^22^. This allows the CBOW model to predict the next word in the sentence given the target word’s neighbouring context words. The trained word representations/embeddings were extracted and integrated into the RNN classifier model.

### Bi-directional LSTM RNN classifier model

The classifier model’s architecture is that of a bidirectional LSTM network, which is a type of Recurrent Neural Network (RNN) and has the advantage of capturing long-term contextual information and is suitable for NLP tasks involving long sequence of words. Similar models have been used for biomedical named entity recognition (BNER) to recognize proteins and gene names and for Relation Extraction tasks from unstructured text^23^. However, relation extraction in the biomedical domain has been mainly restricted to binary relations between entities, such as protein-protein interaction^24^, drug-drug interaction^25^. Here, we evaluate the accuracy of the model to extract relations from three entities incorporating both biological information and experimental methodology into the prediction model. The model takes annotated sentences as inputs and reads it word by word to determine a binary classification of a “yes” or a “no” (1 or 0) if a sentence contains a valid entity relationship between the protein, methodology and subcellular location. The input data was split into a ratio of 1:4, 20% for testing and 80% for training the classifier, making sure the data was separated so that papers did not overlap between the testing and training sets. This prevents the classifier model from analysing very similar input sentences both within the training and testing data sets.

We achieved an average accuracy score of 89.3% (n = 30 experimental runs, standard deviation = 0.028) for all test input sentences in the SUBA testing data set (Table 3, Supplemental table S3-4) (Fig. 3). Interestingly, the classifier was able to provide more accurate predictions for negative input sentences in which the entities did not satisfy semantic and syntactic criteria (95.5%) compared to positive input statements (86.5%). Furthermore, when comparing input sentences with one or more triplet entity groups, the prediction accuracy was observed to be higher for single triplet groups (95.6%), containing a single protein, a subcellular location and an experimental methodology compared to sentences with multiple triplet groups (90.8%). Nevertheless, an overall accuracy of 89.3% is still, to the best of our knowledge, a significant improvement among similar neural network and machine learning approaches used previously for predicting relationships for triplet entity groups using unstructured biomedical text. When evaluating drug-gene-mutation interactions, a graph-based LSTM model achieved an average accuracy of 80.7% for triplet entity relation extraction^26^. Although our model improved accuracy by 8.6% compared to this study, the different dataset, triplet entity group and annotation used in this study make any direct comparison difficult. A comparable study using a distant supervised learning approach for predicting relation between a protein and a subcellular location, achieved an accuracy of 82%^27^, a drop in 7.3% compared with our model and crucially focussed only on binary relations without including experimental methodology as we did in our study.

**Figure 3.**
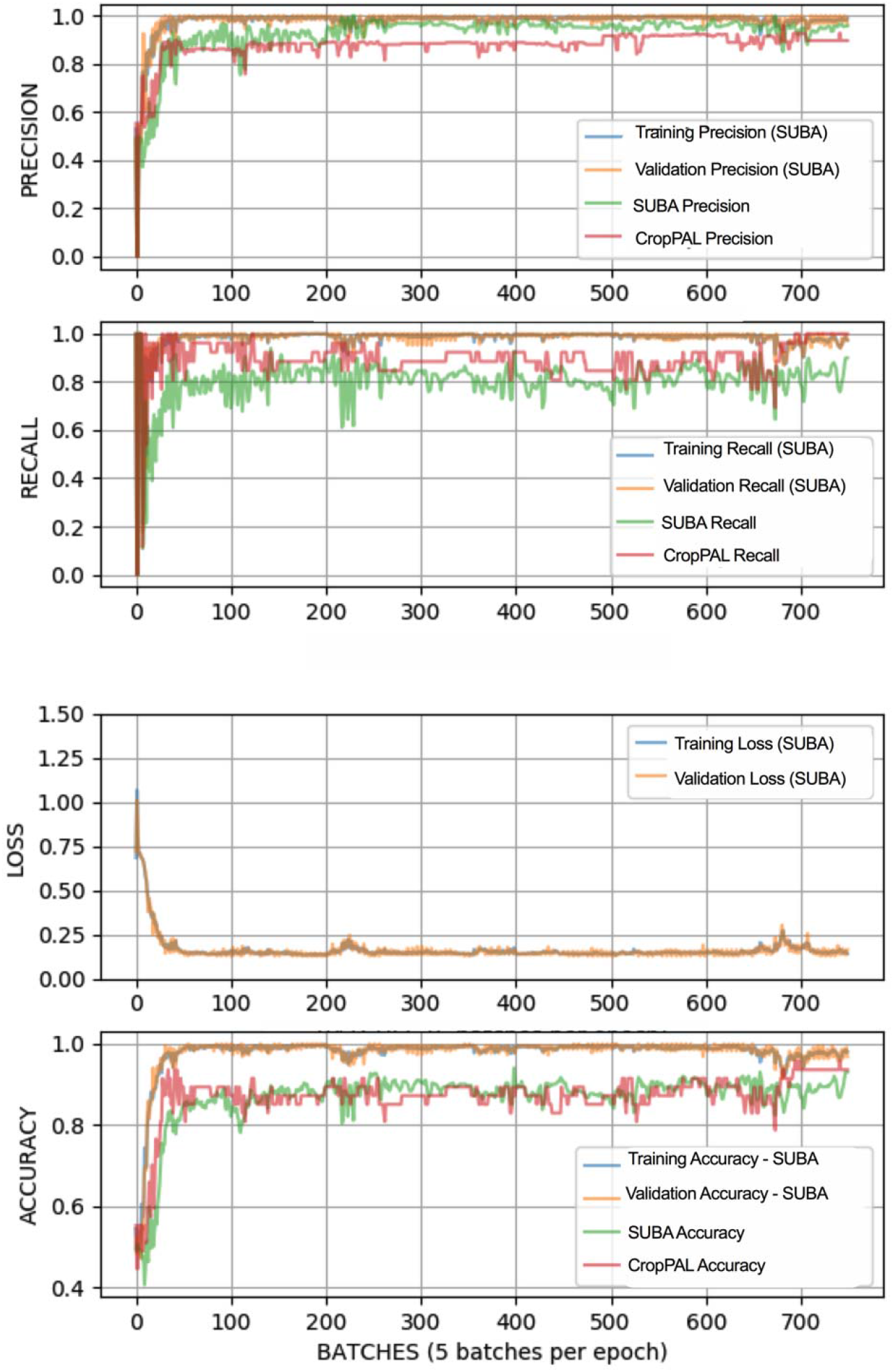
Precision, recall, loss and accuracy progression during the training for the Bi-LSTM to classify entity relationships between proteins, experimental methodology and subcellular locations from input sentences. Training, validation and testing of the model was undertaken using the SUBA dataset and independently tested using the CropPAL dataset.

### CropPAL performance

We next tested the Bi-LSTM RNN for its accuracy in predicting protein subcellular localisation using a dataset independent of the SUBA corpus. The CropPAL (ver 1.0) (http://crop-pal.org/) database stores protein subcellular information for the crop species rice, wheat, barley and maize that is manually curated from the primary literature. Literature retrieval and pre-processing of the paper and input statements were carried out as described for the SUBA dataset (Fig. 1). Using a smaller subset of randomly selected papers from the CropPAL dataset, 65 input sentences were extracted, annotated and tagged using the same process described above. Using the Bi-LSTM classifier, we achieved a similar accuracy score of 0.897 (n = 30 experimental runs, standard deviation = 0.037) as the SUBA dataset demonstrating transfer learning capacity of the model (Table 1, Fig. 3).

**Table 1.** Mean and standard deviation of the precision, recall, accuracy and F1-score for the tested Uni- and Bi-directional LSTM models on the SUBA and CropPAL datasets.

| Dataset | LSTM model | Precision (%) | Recall (%) | Accuracy (%) | F-score (%) |
| --- | --- | --- | --- | --- | --- |
| SUBA | Bidirectional | 95.17 ± 2.4 | 82.80 ± 6.1 | 89.39 ± 2.8 | 88.40 ± 3.6 |
| | Unidirectional | 95.32 ± 1.9 | 79.26 ± 7.1 | 87.80 ± 3.0 | 86.35 ± 4.0 |
| CropPAL | Bidirectional | 89.44 ± 2.6 | 92.56 ± 6.7 | 89.79 ± 3.7 | 90.82 ± 3.8 |
| | Unidirectional | 89.20 ± 2.0 | 89.87 ± 5.6 | 88.37 ± 3.2 | 89.53 ± 3.2 |

### Bi-directional versus Uni-directional LSTM

As a comparison to the bidirectional LSTM, we tested the performance of a unidirectional LSTM (Uni-LSTM) on both SUBA and CropPAL datasets (Table 1). We found the performance dropped marginally for the SUBA dataset, with an average accuracy score of 0.878, a decrease of 1.5% compared to Bi-LSTM. For CropPAL dataset, the accuracy score was also marginally lower, 0.883, a decrease of 1.4% compared to Bi-LSTM. Overall, our analysis suggests that Bi-directional LSTM classifier offers a small improvement in predicting protein subcellular localisation over a uni-directional classifier. However, these results may depend on the type of data used in this study and future analysis will require testing of input sentences that describe other protein characteristics such as protein-protein interaction or protein-protein-disease/trait interaction to analyse the RNN classifier that offers the better overall predictive ability for biological entity relationship extraction.

## Conclusion

We demonstrate the BI-LSTM RNN classifier model uses contextual information in the input sentence to predict triplet entity relationships with a high degree of accuracy and can distinguish from input sentences in which the three entities co-occur by chance. This approach can be effective for extracting biological information from unstructured text in a meaningful way that can be made accessible to the research community. Earlier reports of Machine Learning and neural network models have primarily focussed on amino acid sequence information for predicting protein subcellular localisation with varying degrees of success^28–30^. While these approaches are important for uncharacterised proteins and those predicted from large-scale sequencing datasets, there is a wealth of published protein subcellular information that can be integrated and exploited for practical use by researchers. A Natural Language Processing system that relies on deep learning Technique such as the one described here will help bridge the gap between the data generation process and integrating the vast amount of published molecular and physiological information available about proteins. In particular, a future opportunity exists in improving the sustainability of biological databases by augmenting manual curation efforts with semi or fully automated neural network approaches. Despite their importance in primary research, biological databases face a survivability challenge due to funding cycles typically lasting 3-5 years and essential services such as data curation and maintenance becoming increasingly difficult to sustain^31^. This is best illustrated with the number of biological databases that are listed as obsolete in the annual database issues published by Nucleic Acid Research (NAR) journal. In 2020, NAR reported 125 discontinued databases, with the number of obsolete databases increasing five times over the last five years. In contrast, 65 new database resources were published in the same database issue, with the number remaining steady over the same five year period^32^. While the study here focuses on protein subcellular information, the pipeline developed provides a framework for capturing key functional properties of proteins from published studies and assisting in biocuration efforts. In addition to improving the overall prediction accuracy, we envisage the Bi-LSTM RNN-based pipeline to be applied for extracting relationships involving two or more entities that describe complex protein features such as protein interaction with other biomolecules such as proteins, DNA and RNA as well as for protein-phenotype associations. The transition from manual to semi-automated to potentially, fully automated biocuration systems that rely on deep-learning processes in the future can be exploited by researchers in expanding proteome knowledgebases and for predicting novel functions for individual proteins.

## Methods

### Bulk retrieval of full-text articles and RNN data split

As SUBA does not store full-text articles, we used PubMed ID (or DOI, when PubMed ID was not available) information available from the SUBA stats page (http://suba.live/stats.html) to retrieve full-text articles from various sources. 178 articles were retrieved from the Europe PMC repository open access full-text RESTful API service (https://europepmc.org/RestfulWebService). In addition, full-text articles from the SUBA dataset were also retrieved from the Elsevier text mining service and the CrossRef association of scholarly publishers bringing the total number of articles to 227. The remaining full-text journal articles were retrieved and processed using an in-house developed tool called ‘NLP-ready’ bringing the total number included in the analysis to 1700 published studies. Details for the automated retrieval and processing tool can be accessed from: https://github.com/arabidopsis/NLP-ready

A smaller subset of 200 randomly selected articles from the SUBA corpus were used for the annotation of entity types and classifying entity relationships. The annotated data from this group was used for training and testing the Bi-LSTM recursive neural network classifier. In addition, all 1700 articles were also pre-processed by passing the full text through a Continuous Bag of Words (CBOW) artificial neural network model from Word2Vec^33^ for pretraining the vocabulary as an additional input for the RNN training.

### BRAT annotation

Protein subcellular information and associated methodology in the text was manually annotated using BRAT^16^, using standoff format in which annotation information is kept separate from the text (Fig. 2, Supplemental Fig. S1). Each annotation between a protein, a subcellular location and a methodology include pairwise relations specified through the type of relation and the two entities (Table 2).

**Table 2:** BRAT annotation entity and relationship types defined for the SUBA and CropPAL corpus

| Entity 1 | Entity 2 | Relationship type |
| --- | --- | --- |
| Protein | Subcellular_Location | Located_In |
| Tagged_Protein | Subcellular_Location | Located_In |
| Protein | Methodology | Subcellular_methodology |
| Protein_Locus | Subcellular_Location | Located_In |
| Protein_Locus | Methodology | Subcellular_methodology |
| Protein_variant | Subcellular_Location | Located_In |
| Protein_variant | Methodology | Subcellular_methodology |
| Protein | Identifier | Protein_identifier |
| Protein | Gene_Locus | Locus_link |
| Protein | Subcellular_Location | Not_Located_In |
| Tagged_Protein | Subcellular_Location | Not_Located_In |

### Bi-LSTM model

We propose a bidirectional LSTM model (Fig. 4) that contains an Input layer, Embedding layer, Bi-directional LSTM layer, Hidden layer and an Output layer (Fig. 4)

**Figure 4.**
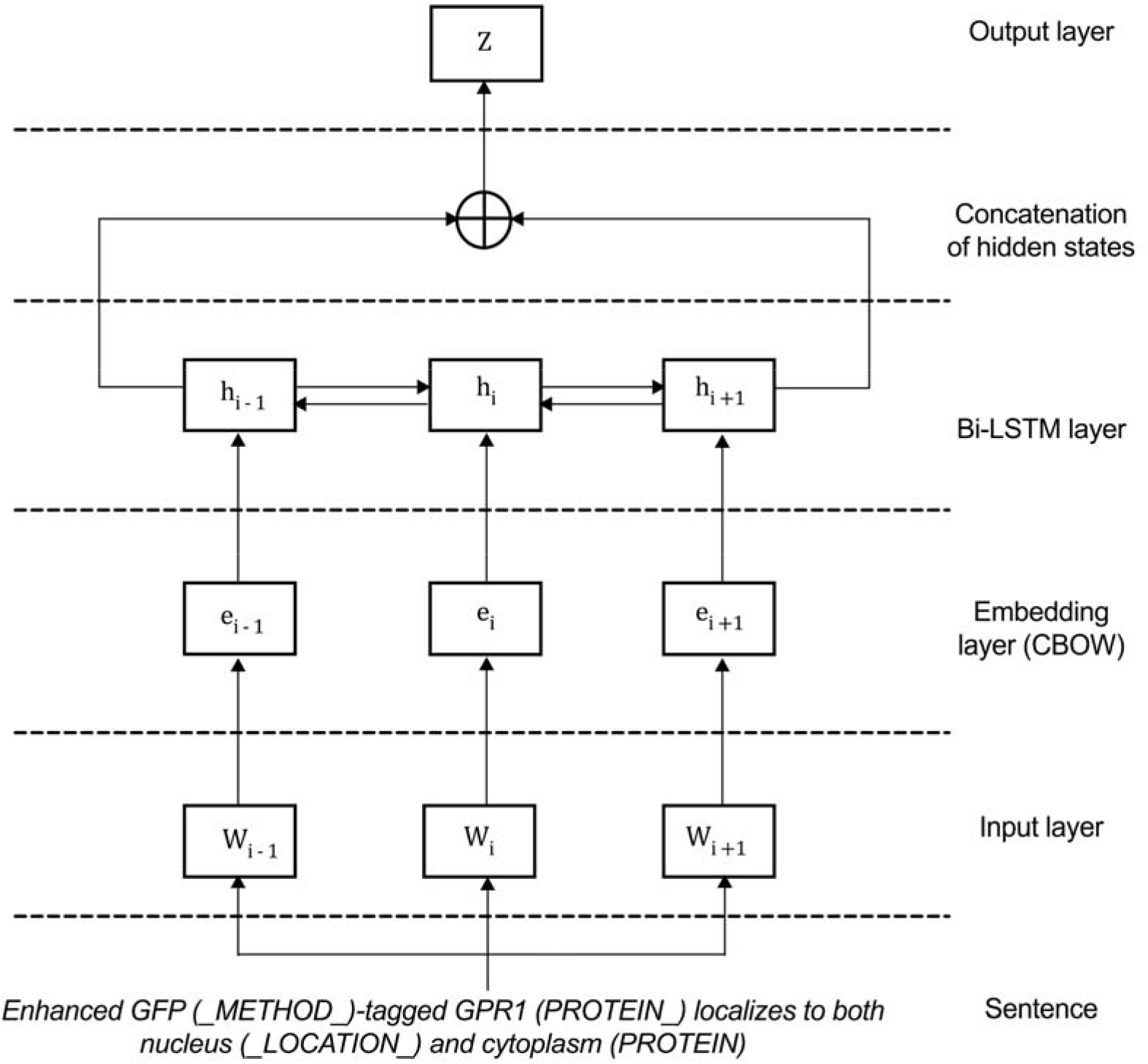
The bi-directional LSTM model to recognize protein subcellular localisation from published literature.

A key limitation of unidirectional LSTMs is that they ignore future context. When reading sequences, it is helpful to get this context of words before you encounter them. To overcome this limitation, in this study we compared both unidirectional and bidirectional LSTMs where one travels in the opposite direction of the other and the hidden states of both are combined at the end:

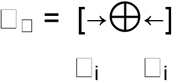

We use concatenation to merge both hidden states and stack 2 layers (4 LSTMs) together and add drop out as regularization.

### Output layer

The output layer contains a dense layer with a Tanh activation. This is then passed through a log softmax and then an argmax which gave us the binary classification:

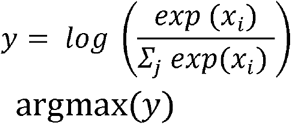

### Experiments

On the training set we performed k-fold cross validation with 5 folds, to achieve the validation and training split ratio of 1:4. When training, the training set was shuffled after each epoch, so that we would be able to perform the k-fold split such that we would not get repeating folds when training. For the 5 folds, each fold was a validation set, making each epoch 5 experiments long.

Model parameters used to generate the results:

Learn rate = 0.01
Dropout rate = 0.50
Hidden nodes = 150
Weight decay = 0.0001
Learn rate decay = 500
Optimizer = Adam

K-folds cross validation was also utilized since there was a sparse amount of data. This was coupled with cross entropy loss to achieve the best results. To test our result we use an F1 score which is the balance between recall and precision.

Where:

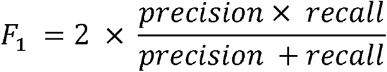

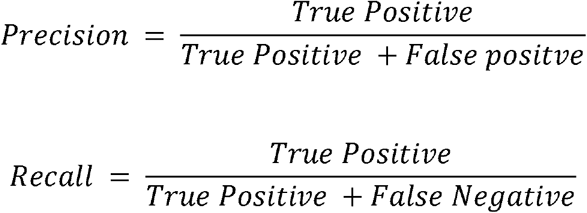

The scores of Recall, Precision, Loss and F1 for 30 experimental runs is provided in the GitHub repository: https://github.com/RhysMenezes/find-a-protein. The output for the Bi-LSTM prediction was independently verified through manual inspection an expert in the field.

## Author Contributions

R.D., G.C., and M.G. designed the study. R.D., J.D.K, R.D.M., I.R.C., and C.M.H. performed the experiments. R.D., J.D.K, and R.D.M. performed data analysis. R.D., J.D.K, R.D.M., G.C., and M.G. prepared and edited the manuscript. All authors approved and edited the final manuscript.

