## Supplemental Figures and Tables for "Identifying protein subcellular localisation in scientific literature using bidirectional deep recurrent neural network"

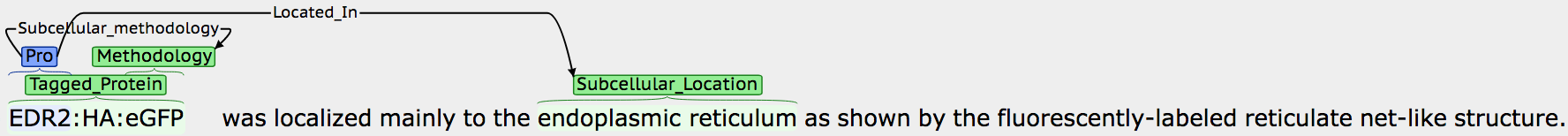

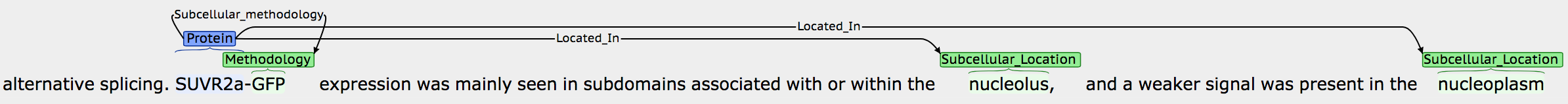

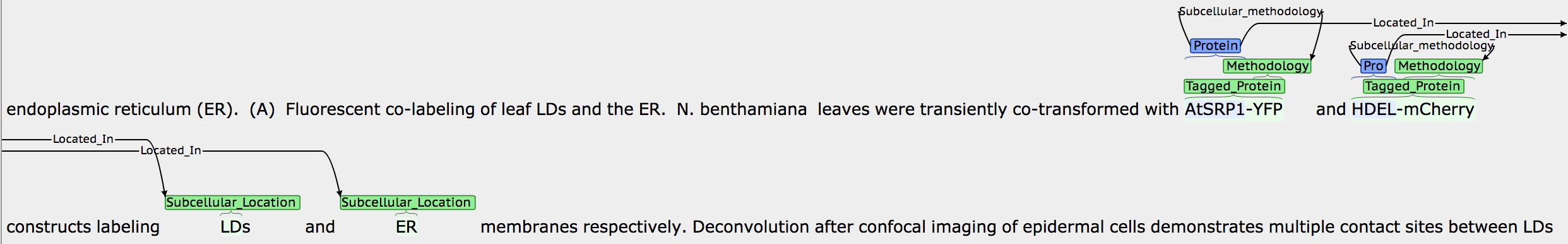

Supplemental Figure 1: Examples of BRAT annotation using extracted text from the SUBA corpus. (A) Triplet entity group representing a tagged protein (concatenation of protein and methodology) localised to a single subcellular component. (B) Triplet entity group containing a tagged protein and two subcellular components. (C) Two triplet groups containing two tagged proteins localised to distinct subcellular compartments.

Table S1: Names derived from the GO subcellular components annotations used in the form of regular expression terms to identify subcellular locations together with co-occurring proteins and experimental methodologies in the SUBA and CropPAL corpus.

| **Major compartments** | **Child location** | **Grandchild locations** |
| --- | --- | --- |
| cytoskeleton (GO:0005856) | intermediate filament  (GO:0005882)  microtubules (GO:0005884)  actin filament (GO:0005884) |  |
| cytosol (GO:0005829) | cell plate (GO:0009504)  cytosolic ribosomes  (GO:0022626) |  |
| endoplasmic reticulum (GO:0005783) | ER lumen (GO:0005788)  ER membrane (GO:0005789) |  |
| extracellular (GO:0005576) | apoplast (GO:0048046)  cell wall (GO:0009505) |  |
| Golgi | Golgi apparatus (GO:0005794) trans-Golgi network (GO:0005802) | Golgi lumen (GO:0005796) Golgi membrane (GO:0000139) multivesicular body (GO:0005771) early endosome (GO:0005769) |
| mitochondrion (GO:0005739) | mitochondrial envelope (GO:0005740) | mitochondrial inner membrane (GO:0005743) mitochondrial outer membrane (GO:0005741) |
|  | mitochondrial matrix  (GO:0005759) |  |
| nucleus (GO:0005634) | nuclear envelope (GO:0005635) | nuclear inner membrane (GO:0005637) nuclear outer membrane (GO:0005640) |
|  | nuclear matrix (GO:0016363) | chromatin (GO:0000785) nucleolus (GO:0005730) |
| peroxisome (GO:0005777) | peroxisomal membrane  (GO:0005778) |  |
|  | peroxisome matrix  (GO:0005782) |  |
| plasma membrane  (GO:0005886) |  |  |
| plastid (GO:0009536) | plastid envelope (GO:0009526) plastid stroma (GO:0009532) plastid thylakoid (GO:0031976) | plastid inner membrane (GO:0009528) plastid outer membrane (GO:0009527) plastoglobules (GO:0010287) plastid thylakoid lumen (GO:0031978) plastid thylakoid membrane (GO:0055035) |
| vacuole (GO:0000325) | vacuole membrane  (GO:0009705) vacuole lumen  (GO:0000330) |  |

Table S2: Regular expression terms to identify experimental methodologies for protein subcellular location data from the SUBA and CropPAL corpus.

| Experimental Methodology | Regular expression query terms |
| --- | --- |
| GFP | “Green fluorescent protein” OR GFP OR FP OR fluorescence OR “fluorescent protein” |
| Mass-spectrometry | MS/MS OR MS OR spectrometry OR “mass spectrometry” |

Table S3: Bidirectional LSTM scores for SUBA and CropPAL datasets

|  | **SUBA** | | | | **CropPAL** | | | |
| --- | --- | --- | --- | --- | --- | --- | --- | --- |
|  | **Accuracy** | **Precision** | **Recall** | **F1** | **Accuracy** | **Precision** | **Recall** | **F1** |
| Run1 | 0.9007 | 0.9407 | 0.8523 | 0.8944 | 0.9362 | 0.9259 | 0.9615 | 0.9434 |
| Run2 | 0.7914 | 0.9479 | 0.6107 | 0.7429 | 0.7872 | 0.9444 | 0.6538 | 0.7727 |
| Run3 | 0.8377 | 0.8906 | 0.7651 | 0.8231 | 0.8936 | 0.8889 | 0.9231 | 0.9057 |
| Run4 | 0.8709 | 0.9661 | 0.7651 | 0.8539 | 0.8936 | 0.8889 | 0.9231 | 0.9057 |
| Run5 | 0.8709 | 0.9825 | 0.7517 | 0.8517 | 0.9362 | 0.8966 | 1.0000 | 0.9455 |
| Run6 | 0.8742 | 0.8881 | 0.8523 | 0.8699 | 0.8723 | 0.8571 | 0.9231 | 0.8889 |
| Run7 | 0.8742 | 0.9826 | 0.7584 | 0.8561 | 0.8936 | 0.8889 | 0.9231 | 0.9057 |
| Run8 | 0.8742 | 0.9826 | 0.7584 | 0.8561 | 0.8511 | 0.8519 | 0.8846 | 0.8679 |
| Run9 | 0.8808 | 0.9520 | 0.7987 | 0.8686 | 0.8723 | 0.8846 | 0.8846 | 0.8846 |
| Run10 | 0.8841 | 0.9831 | 0.7785 | 0.8689 | 0.8936 | 0.8889 | 0.9231 | 0.9057 |
| Run11 | 0.8874 | 0.9197 | 0.8456 | 0.8811 | 0.9362 | 0.8966 | 1.0000 | 0.9455 |
| Run12 | 0.8874 | 0.9389 | 0.8255 | 0.8786 | 0.9149 | 0.8929 | 0.9615 | 0.9259 |
| Run13 | 0.8907 | 0.9462 | 0.8255 | 0.8817 | 0.9149 | 0.8929 | 0.9615 | 0.9259 |
| Run14 | 0.8907 | 0.9531 | 0.8188 | 0.8809 | 0.9362 | 0.9259 | 0.9615 | 0.9434 |
| Run15 | 0.8940 | 0.9209 | 0.8591 | 0.8889 | 0.8723 | 0.8846 | 0.8846 | 0.8846 |
| Run16 | 0.8940 | 0.9916 | 0.7919 | 0.8806 | 0.8511 | 0.8800 | 0.8462 | 0.8627 |
| Run17 | 0.9007 | 0.9760 | 0.8188 | 0.8905 | 0.8723 | 0.9167 | 0.8462 | 0.8800 |
| Run18 | 0.9040 | 0.9348 | 0.8658 | 0.8990 | 0.8723 | 0.8333 | 0.9615 | 0.8929 |
| Run19 | 0.9040 | 0.9688 | 0.8322 | 0.8953 | 0.8723 | 0.8571 | 0.9231 | 0.8889 |
| Run20 | 0.9106 | 0.9552 | 0.8591 | 0.9046 | 0.8511 | 0.8800 | 0.8462 | 0.8627 |
| Run21 | 0.9106 | 0.9621 | 0.8523 | 0.9039 | 0.8723 | 0.8571 | 0.9231 | 0.8889 |
| Run22 | 0.9139 | 0.9424 | 0.8792 | 0.9097 | 0.9574 | 0.9286 | 1.0000 | 0.9630 |
| Run23 | 0.9172 | 0.9429 | 0.8859 | 0.9135 | 0.9362 | 0.9259 | 0.9615 | 0.9434 |
| Run24 | 0.9172 | 0.9493 | 0.8792 | 0.9129 | 0.9362 | 0.9259 | 0.9615 | 0.9434 |
| Run25 | 0.9172 | 0.9559 | 0.8725 | 0.9123 | 0.9362 | 0.9259 | 0.9615 | 0.9434 |
| Run26 | 0.9205 | 0.9562 | 0.8792 | 0.9161 | 0.9149 | 0.8929 | 0.9615 | 0.9259 |
| Run27 | 0.9205 | 0.9562 | 0.8792 | 0.9161 | 0.8936 | 0.8889 | 0.9231 | 0.9057 |
| Run28 | 0.9205 | 0.9630 | 0.8725 | 0.9155 | 0.9362 | 0.9259 | 0.9615 | 0.9434 |
| Run29 | 0.9272 | 0.9441 | 0.9060 | 0.9247 | 0.8936 | 0.8889 | 0.9231 | 0.9057 |
| Run30 | 0.9305 | 0.9571 | 0.8993 | 0.9273 | 0.9362 | 0.8966 | 1.0000 | 0.9455 |
| **Average** | **0.8939** | **0.9517** | **0.8280** | **0.8840** | **0.8979** | **0.8944** | **0.9256** | **0.9082** |
| Stdev | 0.0286 | 0.0245 | 0.0615 | 0.0367 | 0.0376 | 0.0267 | 0.0677 | 0.0385 |

Table S4: Unidirectional LSTM scores for SUBA and CropPAL datasets

|  | **SUBA** | | | | **CropPAL** | | | |
| --- | --- | --- | --- | --- | --- | --- | --- | --- |
|  | **Accuracy** | **Precision** | **Recall** | **F1** | **Accuracy** | **Precision** | **Recall** | **F1** |
| Run1 | 0.8046 | 0.9500 | 0.6376 | 0.7631 | 0.9149 | 0.9231 | 0.9231 | 0.9231 |
| Run2 | 0.8212 | 0.9612 | 0.6644 | 0.7857 | 0.8511 | 0.8519 | 0.8846 | 0.8679 |
| Run3 | 0.8377 | 0.9237 | 0.7315 | 0.8165 | 0.8723 | 0.8846 | 0.8846 | 0.8846 |
| Run4 | 0.8377 | 0.9808 | 0.6846 | 0.8063 | 0.8298 | 0.8750 | 0.8077 | 0.8400 |
| Run5 | 0.8543 | 0.9730 | 0.7248 | 0.8308 | 0.8723 | 0.9167 | 0.8462 | 0.8800 |
| Run6 | 0.8543 | 0.9817 | 0.7181 | 0.8295 | 0.8511 | 0.8800 | 0.8462 | 0.8627 |
| Run7 | 0.8576 | 0.9492 | 0.7517 | 0.8390 | 0.8723 | 0.8846 | 0.8846 | 0.8846 |
| Run8 | 0.8576 | 0.9732 | 0.7315 | 0.8352 | 0.9149 | 0.8929 | 0.9615 | 0.9259 |
| Run9 | 0.8609 | 0.9421 | 0.7651 | 0.8444 | 0.8936 | 0.8889 | 0.9231 | 0.9057 |
| Run10 | 0.8642 | 0.9355 | 0.7785 | 0.8498 | 0.9149 | 0.8667 | 1.0000 | 0.9286 |
| Run11 | 0.8642 | 0.9576 | 0.7584 | 0.8464 | 0.8723 | 0.8846 | 0.8846 | 0.8846 |
| Run12 | 0.8675 | 0.9739 | 0.7517 | 0.8485 | 0.8085 | 0.8696 | 0.7692 | 0.8163 |
| Run13 | 0.8709 | 0.9583 | 0.7718 | 0.8550 | 0.8936 | 0.8889 | 0.9231 | 0.9057 |
| Run14 | 0.8709 | 0.9661 | 0.7651 | 0.8539 | 0.8723 | 0.8846 | 0.8846 | 0.8846 |
| Run15 | 0.8742 | 0.9440 | 0.7919 | 0.8613 | 0.8511 | 0.8800 | 0.8462 | 0.8627 |
| Run16 | 0.8742 | 0.9587 | 0.7785 | 0.8593 | 0.9149 | 0.9231 | 0.9231 | 0.9231 |
| Run17 | 0.8808 | 0.9520 | 0.7987 | 0.8686 | 0.8298 | 0.8750 | 0.8077 | 0.8400 |
| Run18 | 0.8874 | 0.9389 | 0.8255 | 0.8786 | 0.9362 | 0.9259 | 0.9615 | 0.9434 |
| Run19 | 0.8940 | 0.9535 | 0.8255 | 0.8849 | 0.9149 | 0.8929 | 0.9615 | 0.9259 |
| Run20 | 0.8940 | 0.9680 | 0.8121 | 0.8832 | 0.8936 | 0.8889 | 0.9231 | 0.9057 |
| Run21 | 0.8974 | 0.9155 | 0.8725 | 0.8935 | 0.8511 | 0.8800 | 0.8462 | 0.8627 |
| Run22 | 0.9007 | 0.9343 | 0.8591 | 0.8951 | 0.9149 | 0.8667 | 1.0000 | 0.9286 |
| Run23 | 0.9007 | 0.9542 | 0.8389 | 0.8929 | 0.8723 | 0.9167 | 0.8462 | 0.8800 |
| Run24 | 0.9073 | 0.9007 | 0.9128 | 0.9067 | 0.9149 | 0.9231 | 0.9231 | 0.9231 |
| Run25 | 0.9106 | 0.9552 | 0.8591 | 0.9046 | 0.8723 | 0.8846 | 0.8846 | 0.8846 |
| Run26 | 0.9106 | 0.9766 | 0.8389 | 0.9025 | 0.9362 | 0.9259 | 0.9615 | 0.9434 |
| Run27 | 0.9172 | 0.9429 | 0.8859 | 0.9135 | 0.8723 | 0.8846 | 0.8846 | 0.8846 |
| Run28 | 0.9172 | 0.9697 | 0.8591 | 0.9110 | 0.8723 | 0.8846 | 0.8846 | 0.8846 |
| Run29 | 0.9205 | 0.9562 | 0.8792 | 0.9161 | 0.9149 | 0.9231 | 0.9231 | 0.9231 |
| Run30 | 0.9305 | 0.9507 | 0.9060 | 0.9278 | 0.9149 | 0.8929 | 0.9615 | 0.9259 |
| **Average** | **0.8780** | **0.9532** | **0.7926** | **0.8635** | **0.8837** | **0.8920** | **0.8987** | **0.8953** |
| Stdev | 0.0307 | 0.0190 | 0.0711 | 0.0403 | 0.0329 | 0.0205 | 0.0567 | 0.0325 |
